# Sex difference in rat ventilatory response and pulmonary vascular resistance

**DOI:** 10.64898/2026.01.08.697724

**Authors:** L. Ratelet, M. Campillo, L. Parra-Sourdeau, R. Abdou, S. Alibrahimi, A. Delaval, N. Doumbia, R. Masson, J. Preud’Homme, M. Remy, D. Benoist, E. Roux

## Abstract

The aim of this study was to investigate possible sex difference in rat ventilation and pulmonary vascular resistance. Experiments were done on 8 week-old 7 Wistar female and 7 male rats. Under anesthesia, the trachea was connected to a spirometer for measurement of the respiration rate (RR) and, normalized to weight, the tidal volume (Vt) and minute ventilation (VE). Responses to hypoxia/hypercapnia and hypoxia/normocapnia were tested by 1 min respiration in a 20 mL-closed circuit without and with CO_2_ trapping. After euthanasia, blood was collected for hematocrit, the pulmonary circulation was perfused on isolated lungs by a physiological solution at 10-40 mmHg pressure to determine the pressure-resistance curve, and the Fulton ratio (right ventricle/left ventricle+Septum weight ratio) was calculated. Data were compared by Mann-Whitney test and, for pressure-resistance curves, by non linear regression and *F* test. Female weight (226±9.6g) was significantly lower than males ones (356±11.7g). In normoxia/normocapnia, RR was 57±4.0 and 59±4.2 min^-1^, Vt/weight was 1.6±0.2 and 1.5±0.2 mL.100g^-1^, and VE/weight was 89±11 and 90±16 mL. min^-1^.100g^-1^ in female and male rats. Acute hypoxia/hypercapnia and hypoxia/normocapnia significantly increased ventilation. Whatever the conditions, no sex difference were found. Total pulmonary vascular resistance curve was significantly higher in males, whereas the Fulton ratio was similar in females (0.29±0.02) and males (0.28±0.03), as was the hematocrit. In conclusion, no sex difference was identified in the ventilatory response of rats. By contrast, pulmonary vascular resistance was significantly higher in males, without right ventricular hypertrophy.

## Introduction

Rats are animals largely used as models of several human respiratory and pulmonary diseases such as chronic obstructive pulmonary disease (COPD) and pulmonary arterial hypertension (PAH). PAH is a chronic disease characterized by increased pulmonary vascular resistance and pulmonary arterial hypertension, leading to right ventricle hypertrophy and eventually right heart failure. Among the several causes of PAH, one is initial pulmonary hypoxia, which may occur in people living at high altitude or in patients suffering COPD leading to hypoxic pulmonary arterial vasoconstriction, inducing increase in pulmonary artery resistance and chronic pulmonary hypertension with vascular and cardiac remodeling, corresponding to PAH group 3 according to the WHO classification (Kovacs et al., 2024). In humans, a large number of studies have evidenced the impact of sex in the occurrence and the development of PAH, with a complex relationship between sexual hormones and PAH, since women have more risk to develop PAH, but better survival than men, a phenomenon known as the “sex paradox” (Lahm et al., 2014; DesJardin et al., 2024). Though sex difference in pulmonary hypertension is largely documented (Martin and Pabelick, 2014; Bousseau et al., 2023; D’Agostino et al., 2023), most studies using rat model of this disease are done in males only (Aravamudhan et al., 2024; Yun et al., 2024; Zhang et al., 2025).

In normal, i.e., non-pathological conditions, sex difference, both anatomical and functional, in the human cardiovascular system is well-known (Zaid et al., 2023). Regarding the lung, several studies have identified difference between men and women in the anatomy of the rib cage and the architecture of the airways, with physiological consequence in respiratory function at rest and in response to exercise (LoMauro and Aliverti, 2018; Molgat-Seon et al., 2018). Regarding the pulmonary vasculature, it has been shown that pulmonary vessel volume was distributed differently in women and men (Wright et al., 2024), and that total pulmonary vascular resistance was higher in women than in men (Sless et al., 2023).

In rats, few studies have been dedicated to the comparison of respiratory and pulmonary vascular functions in normal animals. Comparison between females and males of different rat strains have shown little difference in respiratory function, with similar minute ventilation (Strohl et al., 1997). Some studies have investigated the pulmonary vascular architecture in murine models and modelled their functional impact, but without sex comparison (Molthen et al., 2004). In rats, pulmonary vascular resistance has been investigated on isolated lung in males (Hillyard et al., 1991; Buncha and Bagi, 2023), females (Rieg et al., 2020) or both (Rubini, 2005), but without sex comparison. Influence of sex hormones has been studied on ovariectomized female rats maintained in chronic hypoxia, in which administration of estradiol increased pulmonary arterial pressure (Kovaleva et al., 2013).

In appears then that few studies have investigated the influence of sex in the pulmonary function of non-pathological rats, though there are used as control in murine models of human pathologies. The aim of this study was to identify the possible influence of sex in the ventilatory physiology and the pulmonary vascular resistance in young adult female and male rats.

## Materials and Methods

### Animal models

Experiments were done on 8-week-old 7 female and 7 male Wistar rats, in accordance with National and European Union guidelines for experimental animal use, using procedures approved by the local Ethical Committee and authorized by National institution. Food and water were available ad libitum, with a 12 h dark/light cycle.

### Ventilatory response in normoxic and acute hypoxic/hypercapnic conditions

Rats were weighted, then anesthetized by intraperitoneal injection of ketamine (10mg/100g body weight) and xylazine (1mg/100g body weight). When anesthetized, animals were placed in decubitus position on a heating pad (37°C), and after local anesthesia with lidocaine, a sagittal incision was done in the skin at the neck, then the underlaying tissues were dissected to give access to the trachea. The trachea was incised in its median part between two cartilaginous rings and cannulated with a Teflon catheter (external diameter 2.80 mm), which was connected to a spirometer head (MLT1L, AD Instruments, Oxford, United Kingdom). Spirometric data were recorded using the Powerlab data acquisition hardware and Labchart software (AD Instruments, Oxford, United Kingdom) for measurement of respiration rate (RR), tidal volume (V_t_) and minute ventilation (V_E_). Responses to hypoxia/hypercapnia and hypoxia/normocapnia were tested after 30 sec and 1 min respiration in a 20 mL-closed circuit without and with CO_2_ trapping by soda lime, respectively.

### Pulmonary vascular resistance, Fulton ratio and hematocrit

After spirometric measurements, anesthetized rats were killed by intraperitoneal injection of sodium pentobarbital (400 mg/kg). After euthanasia, a thoracotomy was done and a 1 mL blood sample was collected in standard heparin tube by intracardiac puncture. The left atrium was sectioned for cardiac discharge, then the lung was gently perfused via puncture of the right ventricle and slow infusion of physiological saline solution (PSS) containing 10 IU heparin to flush out blood. The heart-lung block was then removed out and stored at 4°C in PSS for 2 hours before ex vivo experiments. For measurement of the pulmonary vascular resistance, the isolated heart-lung block was maintained in PSS at room temperature (20 °C). After careful dissection, a suture thread was positioned around the main trunk of the pulmonary artery, which was catheterized through an incision made in the upper part of the right ventricle, and the pulmonary artery trunk was then ligated around the catheter (external diameter 1 mm). The catheter was then connected to the perfusion system composed of an open syringe containing PSS and a Teflon tubing of known resistance (*R*_*t*_). The lung vasculature was perfused with PSS at various hydrostatic pressures ranging from 10 mmHg to 40 mmHg. The time needed for 2 mL perfusion was measured at each hydrostatic pressure, allowing the calculation of the perfusion flow rate. Considering that the relationship between flow rate (*J*), hydrostatic pressure of perfusion (*P*), and resistance of the whole circuit (*R*) is given by the following equation:

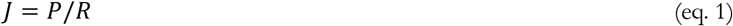

and that the total circuit resistance *R* is the sum of the resistance of the tubing system (*R*_*t*_) and the inherent resistance of the pulmonary vascular network (*R*_*p*_), *R*_*p*_ was therefore calculated using the following equation:

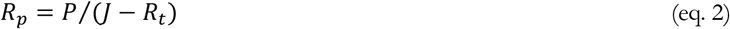

After measurement of the pulmonary perfusion, the heart was removed, the right ventricle and the left ventricle + septum were dissected and weighted, in order to calculate the Fulton ratio, i.e., the weight ratio of the right ventricle on the left ventricule+septum.

Hematocrit was measured in triplicate on heparined blood.

### Statistical analysis

Data are given as mean±SEM. For spirometric measurement, the respiration rate, the tidal volume and the minute ventilation were compared by Mann-Whitney test. Changes in spirometric values induced by 30 sec and 1 min hypoxia/hypercapnia and hypoxia/normocapnia were expressed as percentage of normoxia/normocapnia values and compared by linear regression analysis. Pressure-resistance curves were fitted by nonlinear regression analysis using a first-order exponential decay equation and compared by *F* test. Hematocrit and Fulton ratio were compared by Mann-Whitney test. Differences were considered significant when *p*<0.05.

## Results

### Ventilatory parameters

#### Normoxia

As expected, female weight (226±9.6g) was significantly lower than male ones (356±11.7g), corresponding to 20% difference (*p*=0.0012). In female and male rats, the respiration rate was 57±4.0 and 59±4.2 cycles per min, almost identical (5% difference) and not statistically different (*p*=0.94) (Figure 1A). Tidal volume was 16% lower in females (3.5±0.36 mL) than in males (5.4±0.93mL), but the difference was not significant (*p*=0.06). Minute ventilation was 18% lower in females (197±20 mL.min^-1^) than in males (319±63 mL.min^-1^), without statistical significance (*p*=0.13). When normalized to weight, tidal volume minute ventilation appeared to be almost similar, with 1% and 5% difference between sexes, respectively (Figure1B and C). Individual data are given in Supplemental table 1.

**Figure 1.**
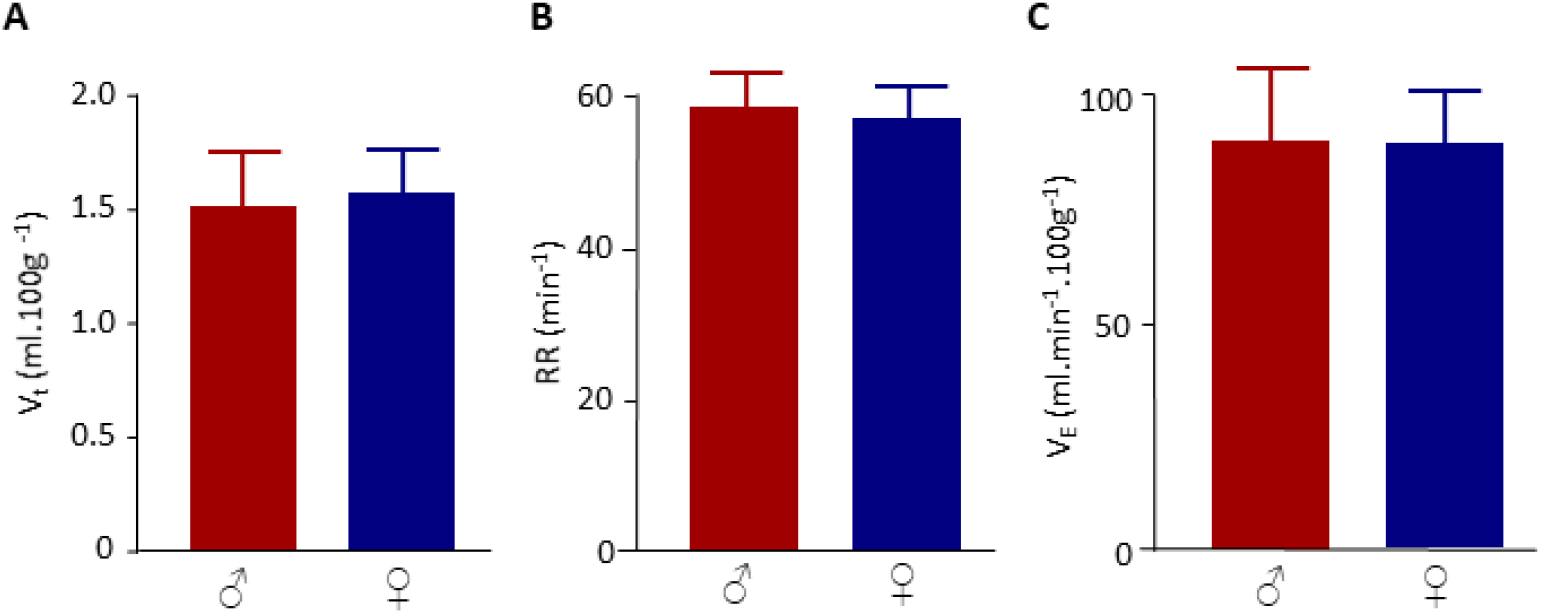
Female and male normoxic ventilation. Red symbols: male. Blue symbols: female. A. Tidal volume (V_t_). B. Respiration rate (RR). C. Minute ventilation (V_E_). Column are mean values; error bars are SEM (n=6). Comparison between females and males were non-significant (Mann-Whitney test, *p*>0.05).

#### Acute hypoxia with and without hypercapnia

When submitted to acute hypoxia without hypercapnia, both male and female rats exhibited an increase in respiration rate, tidal volume and, as a consequence, minute ventilation (Figure 2A, B and C). Linear regression showed no significant difference between female and male response to acute normocapnic hypoxia (Table 1). In acute hypercapnic hypoxia, the ventilatory response was almost similar to that in normocapnic hypoxia, and no statistical difference were identified between female and male responses (Figure 2D, E and F and Table 1). Individual data are given in Supplemental table 2.

**Table 1.**
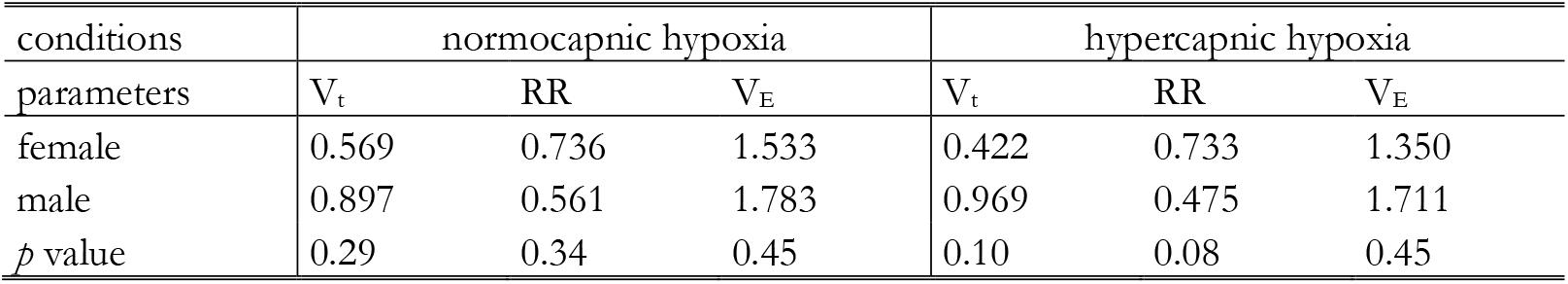
Regression slopes of the ventilatory response to acute normocapnic and hypercapnic hypoxia. Lines 1 and 2: values of the slopes of the linear regression of the increase in tidal volume (Vt), respiration rate (RR) and minute ventilation (V_E_) in females (line 1) and males (line 2), as presented in Figure 2A to F. Line 3: *p* values of the comparison of the linear regression in females and males.

| conditions | normocapnic hypoxia |  |  | hypercapnic hypoxia |  |  |
| --- | --- | --- | --- | --- | --- | --- |
| parameters | $V_t$ | RR | $V_E$ | $V_t$ | RR | $V_E$ |
| female | 0.569 | 0.736 | 1.533 | 0.422 | 0.733 | 1.350 |
| male | 0.897 | 0.561 | 1.783 | 0.969 | 0.475 | 1.711 |
| $p$ value | 0.29 | 0.34 | 0.45 | 0.10 | 0.08 | 0.45 |

**Figure 2.**
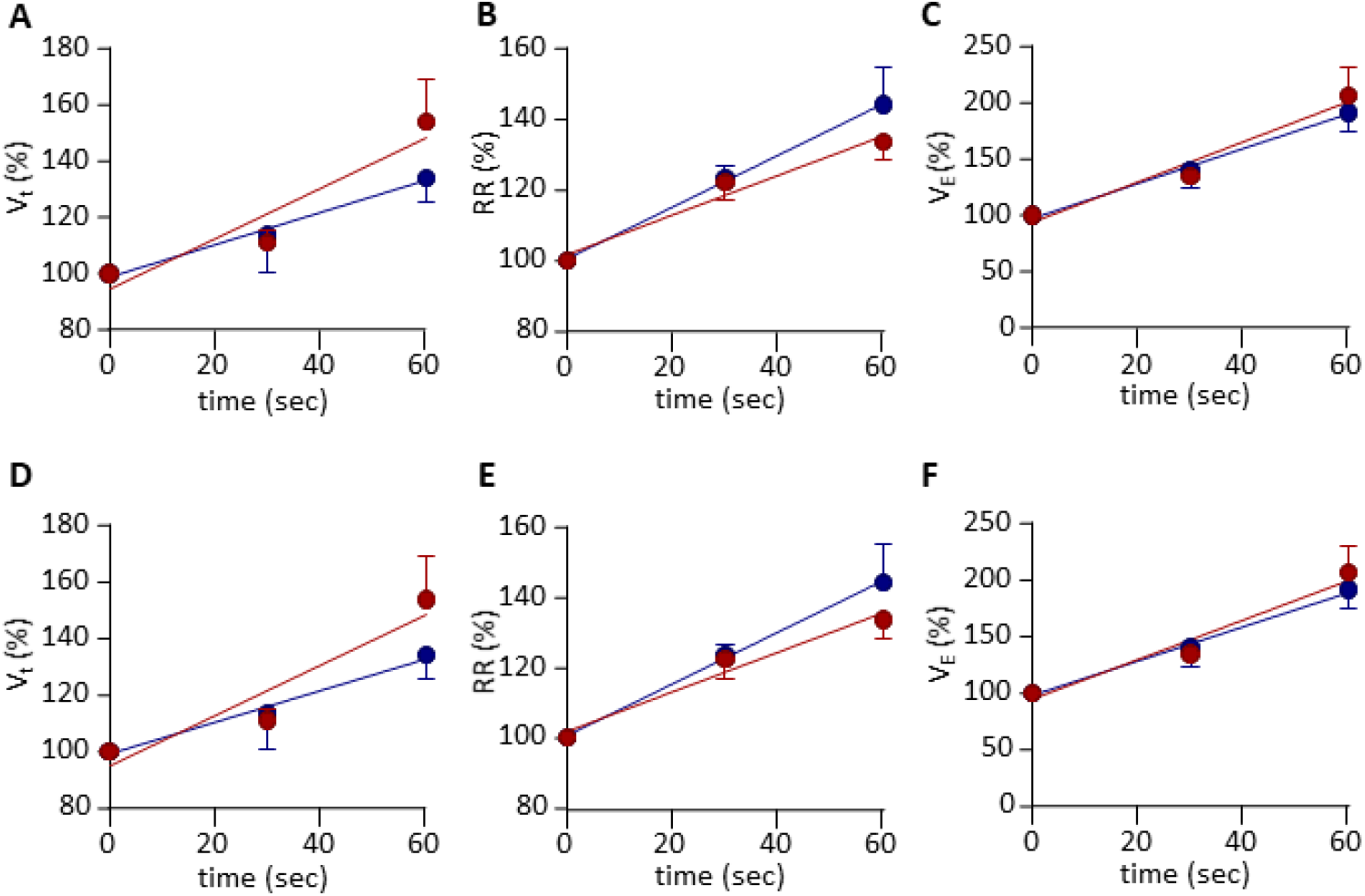
Female and male ventilatory response to acute hypoxia. Red symbols: male. Blue symbols: female. Change in ventilatory parameters of female and male rats, in % of normoxic normocapnic conditions, induced by acute normocapnic (A, B and C) and hypercapnic (D, E and F) hypoxia. Dots are mean values, error bars are SEM (n=6). Vt: tidal volume (A, D). RR: respiration rate (B, E). V_E_: minute ventilation (C, F). Comparison between females and males were non-significant (linear regression analysis, *p*>0.05).

**Figure 3.**
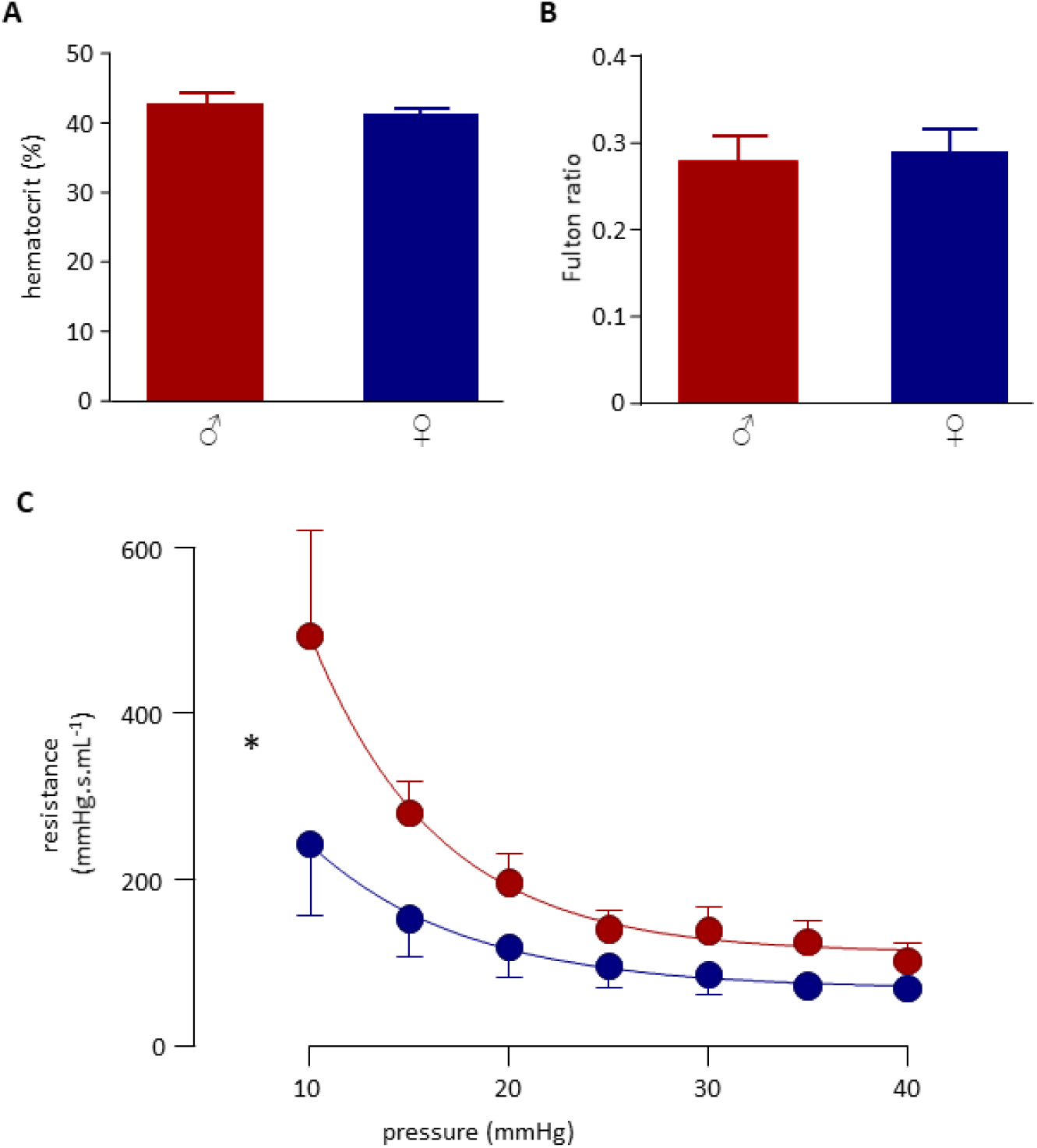
Female and male hematocrit, Fulton ratio and pulmonary vascular resistance. Red symbols: male. Blue symbols: female. A. Hematocrit. B. Fulton ratio (right ventricle/left venticle+septun weight ratio). A, B: column are mean values, error bars are SEM (n=7). Comparison between females and males were non-significant (Mann-Whitney test, *p*>0.05). C. Pulmonary vascular resistance curve. Abscissa: perfusion pressure. Ordinate: pulmonary vascular resistance. Dots are mean values, error bars are SEM (n=5). * *p*<0.05, nonlinear regression analysis, *F* test.

### Hematocrit

Hematocrit values showed no significant difference between males and females (Figure 2A). Taken together, the hematocrit was 42.1±0.88.

### Fulton ratio

Fulton ratio values showed no significant difference between males and females (Figure 2B). Taken together, the Fulton ratio was 0.29±0.018. Individual data of hematocrit and Fulton ratio are given in Supplemental table 3.

### Pulmonary vascular resistance

The values of the pulmonary vascular resistance were fitted by the following first-order exponential decay equation:

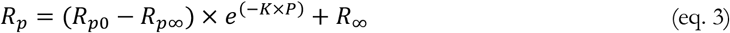

where *P* is the pressure (in mmHg), *R*_*p*_ is the pulmonary vasculature resistance (in mmHg.s.mL^-1^), *R*_*p*0_ is *R*_*p*_ for *P*=0, *R*_*p*∞_ is *R*_*p*_ for *P* = ∞, and *K* is the rate constant (in mmHg^-1^). Whatever the perfusion pressure, the pulmonary vascular resistance was higher in males than in females, and the resistance curves were statistically different (*p*=0.0003) (Figure 2C). Individual data are given in Supplemental table 4.

## Discussion

The results of this study show that ventilation at rest in normoxic conditions did not differ between female and male adult Wistar rats. Acute hypoxia, with and without hypercania, induced in both sexes a time-dependent increase in minute ventilation, due to both increase in tidal volume and respiration rate, without sex difference. Pulmonary vascular resistance was significantly lower in females than in males, whereas the Fulton ratio and the hematocrit were similar in both sexes.

### Ventilation

Average male and female minute ventilation found in our study was higher than that previously found in for Sprague-Dawley, Brown Norway, Zucker and Koletsky rats strains, whereas the respiration rate found in these strains was higher and the tidal volume lower than that of our study (Strohl et al., 1997). These differences are probably due not to difference in rat strains but to experimental conditions for ventilation measurements, i.e., vigil *versus* anesthetized and cannulated animals. The absence of sex difference observed in our study is in accordance with this previous investigation done in other rat strains, were significant difference was observed only in respiration rate, with was higher in females than males, but not in tidal volume and minute ventilation (Strohl et al., 1997). In humans, difference in airways and rib cage anatomy have been showed to be responsible for sex difference in respiratory function, with lower ventilatory values at rest and differences in response to exercise (LoMauro and Aliverti, 2018; Molgat-Seon et al., 2018). Since rat and human anatomy are quite different, extrapolation of anatomical impact on respiratory function from one species to another has limited relevance.

Close-circuit breathing causes progressive hypoxia and hypercapnia, which may activate hypercapnic and/or hypoxic chemoreflex, leading to increased ventilation. As expected, our results show time-dependent increase in ventilation parameters. In our study, response to hypoxia was similar with and without hypercapnia, suggesting that hyperventilation was triggered by only hypoxic, and not hypercapnic, chemoreflex. This may be due to the fact measurements were done during short-time exposure, i.e., 1 minute, whereas the time constant of the central chemoreflex, responsible for 70-80% of the hypercapnic response, is longer, around 60-150 sec (Vassilakopoulos, 2012). Regarding sex difference, during acute hypoxia, with or without hypercapnia, change in minute ventilation was very similar in females and males, a result in accordance with data obtained in other rat strains (Strohl et al., 1997). However, males and females did not seem to respond in an identical manner. Though the difference was not significant, male responded to acute hypoxia with higher tidal volume and lower respiration rate. Increased frequency in females vs males in acute hypoxia or hypercapnia has already been observed in other rat strains (Strohl et al., 1997). During exercise, it has been shown in humans that respiration rate increase was high in women than men. This suggests that sensitivity to hypoxia may differ with sex, even when leading to similar ventilation.

### Pulmonary vasculature

Our experiments on isolated lungs showed a large and significant sex-difference in pulmonary vascular resistance, which was lower in females for all tested perfusion pressures, ranging from 10 to 40 mmHg, i.e., lower and higher than the average normal pulmonary arterial pressure, i.e., 15-20mmHg in rats (Kwan et al., 2024). Since experiments were performed on non-ventilated lungs perfused with non-oxygenated physiological saline solution at room temperature, i.e., far below normal body temperature, the difference in vascular resistance observed between females and males in our study is unlikely to be due to difference in vascular reactivity, an active function that requires normal physiological conditions incompatible with the experimental protocol. More likely, the difference may be due to difference in the anatomical architecture of male and female pulmonary vasculature. Further studies are needed to determine the relative contributions of vascular architecture and vascular reactivity in sex difference in pulmonary vascular resistance. Though sex difference in pulmonary hypertension is largely documented (Martin and Pabelick, 2014; Bousseau et al., 2023; D’Agostino et al., 2023), most studies using rat model of this disease are done in males only (Aravamudhan et al., 2024; Yun et al., 2024; Zhang et al., 2025). Influence of sex hormones has been studied on ovariectomized female rats maintained in chronic hypoxia, in which administration of estradiol increased pulmonary arterial pressure (Kovaleva et al., 2013). Pulmonary vascular resistance has been investigated on isolated lung in males (Hillyard et al., 1991; Buncha and Bagi, 2023), females (Rieg et al., 2020) or both (Rubini, 2005), but without sex comparison, and physiological values were usually not mentioned in these publications, only variations normalized to control values. In Wistar rat ventilated lungs perfused in situ with a plasma-like colloid solution, pulmonary vascular resistance averaged on both sexes was 8 cmH_2_O.min. mL^-1^, corresponding to 350 mmHg.sec. mL^-1^, for a perfusion pressure of 25 mmHg (Rubini, 2005), a value in the same range but higher than our own results. However, considering the different viscosities of the perfusion solutions, i.e., plasmatic *versus* saline physiological solutions, resistance values obtained in this study were quite similar as ours. Since vascular resistance has not been measured with blood, which viscosity is much higher due to blood cells, pulmonary vascular resistance is supposedly much higher in real physiological conditions. However, since hematocrit was similar in males and females, the difference between sexes observed in our study is likely to be similar with blood lung perfusion.

Despite the difference observed in total pulmonary vascular resistance between males and females, the Fulton ratio, an index of right ventricular hypertrophy that is increased in pulmonary hypertension, was similar in both sexes, indicating similar right ventricular development and suggesting close pulmonary blood pressure. These data are in accordance with the literature, showing similar mean pulmonary arterial blood pressure in physiological conditions in male, female and ovariectomized female Sprague-Dawley rats, and similar Fulton ratio values (0.26-0.28) (Kwan et al., 2024), close to our own data. In seems hence that the normal pulmonary cardiopulmonary function, i.e., the ability to ensure lung perfusion in physiological conditions, is similar in both sexes, but not achieved in the same way.

In conclusion, our study showed no sex difference in ventilatory physiology in adult Wistar rats, either in normoxia or acute hypoxia, with and without hypercapnia. By contrast, pulmonary vascular resistance was significantly higher in males, without right ventricular hypertrophy. These sex differences in pulmonary vascular physiology may contribute to the differential susceptibility between males and females to develop pulmonary hypertension.

